# Baseline psychological traits contribute to Lake Louise Acute Mountain Sickness score at high altitude

**DOI:** 10.1101/2021.07.22.451589

**Authors:** Benjamin James Talks, Catherine Campbell, Stephanie J Larcombe, Lucy Marlow, Sarah L Finnegan, Christopher T Lewis, Samuel J E Lucas, Olivia K Harrison, Kyle TS Pattinson

**Affiliations:** Population Health Sciences Institute, Newcastle University, Newcastle Upon Tyne, United Kingdom; Birmingham Medical Research Expeditionary Society, Birmingham, United Kingdom; Medical School, University of Birmingham, Birmingham, United Kingdom; Warwick Medical School, Warwick University, Coventry, United Kingdom; Nuffield Department of Clinical Neurosciences, University of Oxford, United Kingdom; Department of Anaesthesia, Ysbyty Gwynedd, Bangor, United Kingdom; School of Sport, Exercise and Rehabilitation Sciences, University of Birmingham, Birmingham, United Kingdom; Translational Neuromodeling Unit, Institute for Biomedical Engineering, University of Zurich and ETH Zurich, Switzerland; School of Pharmacy, University of Otago, New Zealand

**Author notes:** **Corresponding author** Dr Benjamin James Talks, Newcastle University, Newcastle Upon Tyne, Tyne and Wear, NE1 7RU, United Kingdom.

**Keywords:** interoception, altitude, breathlessness, filter detection task, exercise, acute mountain sickness

## Abstract

**Background:** Interoception refers to an individual’s ability to sense their internal bodily sensations. Acute mountain sickness (AMS) is a common feature of ascent to high altitude that is only partially explained by measures of peripheral physiology. We hypothesised that interoceptive ability may explain the disconnect between measures of physiology and symptom experience in AMS.

**Methods and Material:** Two groups of 18 participants were recruited to complete a respiratory interoceptive task three times at two-week intervals. The control group remained in Birmingham (140m altitude) for all three tests. The altitude group completed test 1 in Birmingham, test 2 the day after arrival at 2624m, and test 3 at 2728m after an 11-day trek at high altitude (up to 4800m).

**Results:** By measuring changes to metacognitive performance, we showed that acute ascent to altitude neither presented an interoceptive challenge, nor acted as interoceptive training. However, AMS symptom burden throughout the trek was found to relate to sea-level measures of anxiety, agoraphobia, and neuroticism.

**Conclusions:** This suggests that the Lake Louise AMS score is not solely a reflection of physiological changes on ascent to high altitude, despite often being used as such by researchers and commercial trekking companies alike.

## Introduction

Acute Mountain Sickness (AMS) is a common feature of ascent to altitude, affecting ~25% of individuals ascending to moderate altitudes (2000-3000m) and up to 58% of individuals at 4500m (Honigman et al., 1993; Schneider et al., 2002). Lake Louise AMS Score, a self-reported symptom score, is the most widely used measure of AMS (Roach et al., 2018). It is well recognized that symptoms of Lake Louise AMS score are not fully explained by measures of hypoxic insult, including arterial oxygen saturations, respiratory rate, and heart rate (Chen et al., 2012; Wagner et al., 2012). Direct measures of hypoxia on brain function also fail to explain this disparity, including matched regional oxygen saturations, electroencephalography, cerebral blood flow velocity, and cerebral oedema (Mairer et al., 2012; Feddersen et al., 2015). Given the diverse array of subjective symptoms induced by ascent to high altitude (Hall et al., 2014), it may be that discrepancies in symptoms reporting between individuals can be explained by differences in their perceptual sensitivity or behavioural profiles.

Interoception refers to an individual’s ability to sense the internal state of their body (Simmons and Land, 1987; Barret and Simmons, 2015). The “Bayesian Brain” hypothesis of perception, including interoception (Barret and Simmons, 2015; Stephan et al., 2016), is a popular neuroscientific theory. In brief, the Bayesian Brain Hypothesis proposes that in order to interpret numerous noisy sensory stimuli (e.g. vision, pain, nausea), the brain generates an internal model of the world, against which it constantly tests new sensory inputs against. The second-order process assessing the accuracy of this predictive model is known as metacognition, a term used to describe “cognition about cognition” (Stephan et al., 2016), or “insight” into your own perceptions. By extension, as all symptoms are produced centrally in the brain, they cannot be fully explained by measures of peripheral physiology alone. Differences in interoception may help explain the discrepancies in AMS symptomology between individuals on ascent to high altitude.

Ascent to high altitude is associated with an array of physiological and behavioural stressors, including the disruption of multiple physiological symptoms due to AMS (Hall et al., 2014), hypoxia and its associated systemic inflammatory response (Eltzschig and Carmeliet, 2011), and fatigue resulting from travel across multiple time zones (Stephan et al., 2016). Such stressors have the potential to impair interceptive performance either independently or in combination. Interestingly, habitual exercise is thought to improve interoceptive performance, with athletes demonstrating better matching between ventilation and perceived breathlessness than sedentary controls (Faull et al., 2016). Further, functional magnetic resonance imaging of brain networks associated with anticipating breathing stimuli has shown brain activity that reflects subsequent interoceptive perceptions in athletes compared to sedentary controls (Faull et al., 2018), and changes in activity in patients with chronic respiratory disease after a course of pulmonary rehabilitation (exercise and education) on exposure to breathlessness cues (Hergistad et al., 2017). Therefore, this study aimed to test the hypothesis that initial ascent to high altitude would impair interoceptive performance, while daily exercise in the form of an 11-day trek at high altitude would act as interoceptive training and thus improve performance.

Individuals’ psychology can also play a significant role in their experience of symptoms – framing their internal model of the world according to the Bayesian Brain Hypothesis. In particular, fatigue and anxiety may be the brain’s manifestations of poor perceived self-efficacy and control, presenting as a state of learned helplessness (Stephan et al., 2016). Indeed, one study of breathlessness in 100 patients with Chronic Obstructive Pulmonary Disease demonstrated different behavioural profiles and brain activity in the anterior insula (a likely key interoceptive centre) between a high and low symptom burden group, in the absence of any differences in spirometry between the two groups (Finnegan et al., 2021). Therefore, we also hypothesised that self-report questionnaires characterising individuals’ baseline psychological state may correlate with their symptom burden over the expedition.

## Methods

### Study Design

This study was a two-group repeated measures design, consisting of equally sized altitude and control groups. Both groups completed interoceptive testing at three time points two weeks apart. The control group completed all three tests in Birmingham, United Kingdom (140m). The altitude group completed baseline testing in Birmingham, United Kingdom, (140m); after arrival at high altitude in Lachung, India (2624m); and after completing a 11-day trek at altitude in Lachen, India (2728m). The ascent profile of the trek is shown on Figure 1, with the highest camp situated at 4800m. This study was approved by the Central University Research Ethics Committee of the University of Oxford (Ref: R60699/RE001). All participants provided written consent.

**Figure 1.**
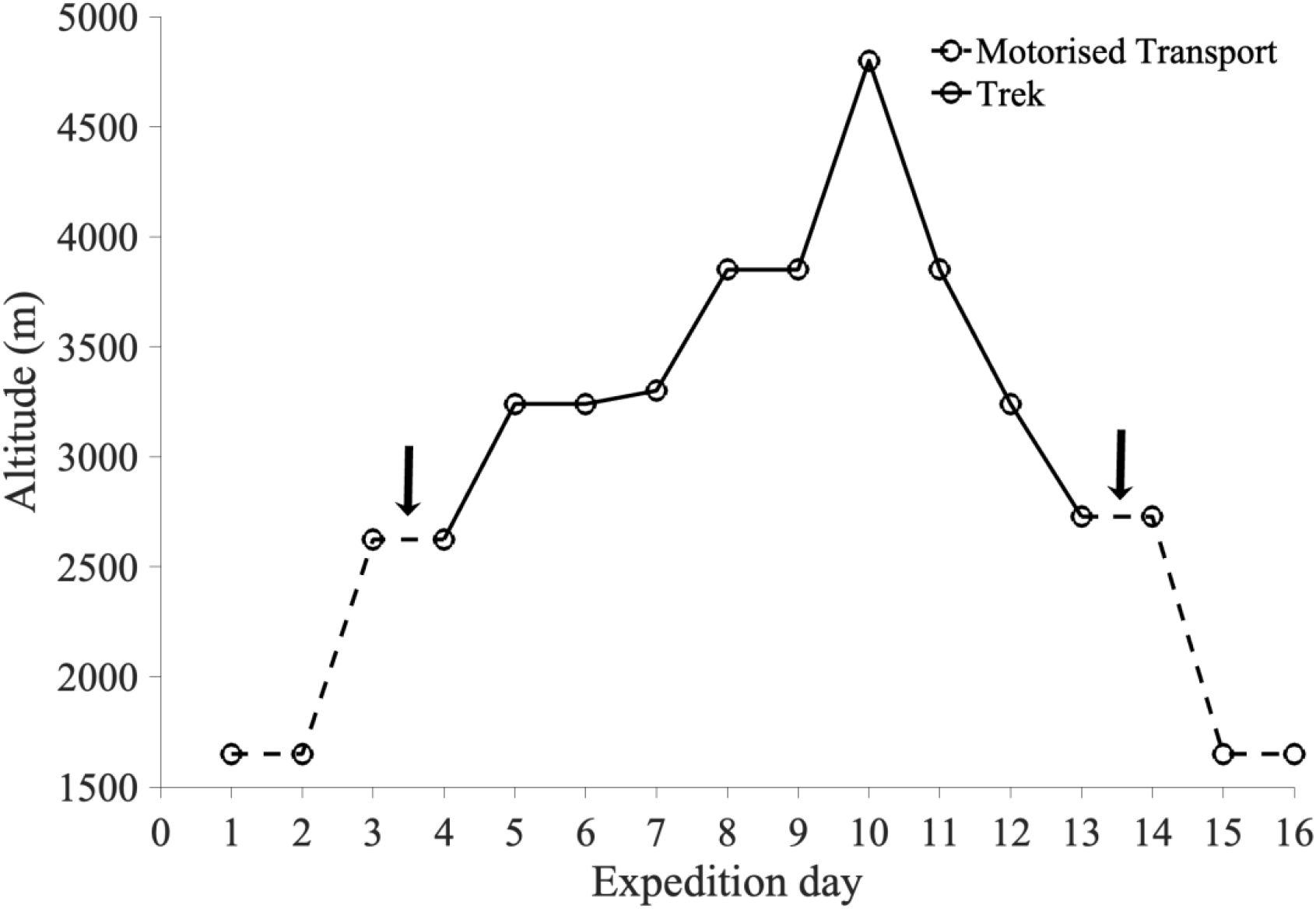
The ascent profile of the altitude group in Sikkim, India. Days of travel by motor vehicle are plotted in a dashed line and days of trekking by foot are plotted with a solid line. The times of the two interoceptive tests on the expedition are marked by arrows.

### Study Participants

The altitude group was composed of members of the Birmingham Medical Research Expeditionary Society (n = 18, 11 males, 7 females, age 23-72 years) taking part in a two week trek in Sikkim, India. An age- and sex- matched control group was recruited through advertisement on the University of Birmingham campus. Exclusion criteria for participating in the study included significant medical comorbidities, smoking history, recent travel across multiple time zones, and recent ascent to high altitude (see Supplemental Material for full criteria). The mean age difference between matched participants was 0.83 years (range 0-7 years).

### Primary Outcome Measures

#### Respiratory Interoceptive Test

A respiratory threshold detection task, the filter detection task (Harrison et al., 2020a), was used as a measure of respiratory interoception. In this task, the participant breathes through a simple breathing system, and following three baseline breaths, either an inspiratory load is added via the addition of clinical breathing filters, or an empty filter (sham) is used. After each trial, participants are asked to decide whether or not resistance was added, as well as reporting their confidence in their decision on a confidence scale from 0 to 10. The number of filters is varied according to an algorithm that tracks performance, until a threshold is found at which the participant is 60-85% confident in their response. The task is then repeated at this threshold for 60 trials. The filter detection task can then be used to determine perceptual sensitivity (number of filters), perceptual bias in symptom reporting (bias towards yes or no), metacognitive bias (average confidence), and metacognitive performance (Mratio, calculated from meta-d’/d) – i.e. the ability to accurately reflect upon and thus control cognitive or perceptual processes (Garfinkel et al., 2016a, 2016b).

#### Cardiorespiratory Physiology

Basic measures of cardiorespiratory physiology were made non-invasively using pulse oximetry and an automatic sphygmomanometer, measuring oxygen saturations, heart rate, and blood pressure. These measures were taken at the time of each filter detection task for both the altitude and control group, and daily in the altitude group during the trek as part of a daily medical review.

#### Self-Report Scores

Participants were asked to complete a number of self-report scores during each of their three testing sessions including the Lake Louise AMS Scale (Roach et al., 2018), Multidimensional Assessment of Interoceptive Awareness (Mehling et al., 2012), Fatigue Severity Scale (Krupp et al., 1989), Epworth Sleepiness Scale (Johns, 1991), State-Trait Anxiety Inventory (Spielberger et al., 1983), Center for Epidemiological Studies Depression Scale (Radloff, 1977), Positive and Negative Affect Scale (Watson et al., 1988), Mobility Inventory for Agoraphobia (Chambless et al., 1985), Anxiety Sensitivity Index (Reiss et al., 1986), UK Biobank Neuroticism Scale (Smith et al., 2013), and Who Global Physical Activity Scale (World Health Organization) reported in metabolic equivalent minutes. The altitude group also completed daily Lake Louise AMS scores throughout the trek.

### Statistical Analysis

Statistical analysis was performed according to the pre-published statistical analysis plan (https://osf.io/zgj9c/).

#### Examining Interoceptive Performance

The respiratory threshold task was analyzed using the hierarchical HMeta-d statistical model (Harrison et al., 2020a); model fits were implemented in MATLAB (Mathworks, Natick, United States of America) with sampling conducted using JAGS (Plummer, 2003). The HMeta-d model was fitted separately for each pair of tests in the control group and altitude group to look at the effect of initial ascent to high altitude (visit 1 and 2) and of an 11-day trek at altitude (visit 2 and 3). Mratio was compared between each visit by calculating a one-tailed 95% Highest Density Interval (HDI) across the distribution of samples for each visit, a significant difference was defined as a HDI not spanning zero. The effect of ascent to high altitude and exercise at altitude were examined by looking at the interaction effect between the control and altitude group for the aforementioned paired timepoints.

Additional variables of the filter detection task, including perceptual sensitivity, perceptual bias, and metacognitive bias, are not fit hierarchically within the Hmeta-d model, so can be compared using standard frequentist statistics. Repeated measures analysis of variance (RANOVA) was used to compare these measures between visits with a 5% significance level. To compare the self-report scores between the altitude and control groups for each visit, responses were first tested for normality using an Anderson-Darling test, then if normal compared using a two-tailed t-test and if non-parametric compared using a Wilcoxon rank sum test. Uncorrected p-values are presented and a Bonferroni correction for multiple comparisons between the two groups has been used to adjust the significance level, p = 0.05/20 = 0.0025.

To investigate the relationship between metacognitive ability and AMS, linear regression models were used to compare metacognitive performance from baseline (visit 1) and arrival at high altitude (visit 2) with AMS symptom burden over the trek. A significant relationship between the covariate and log of Mratio was defined as a two-tailed HDI that does not span zero across the beta samples for the covariate.

#### Examining Baseline Psychology

Hierarchical cluster analysis, similar to the method outlined by Abdallah et al., (2019), was used to assess the relationship between measures of symptoms of AMS on ascent to high altitude and baseline psychological state. Means of the daily measures of Lake Louise AMS score and oxygen saturations from the trek were included to represent AMS symptom burden.

Only self-report scores that referred to individuals’ “usual state” rather than the present moment or preceding week were included in the cluster analysis to best represent their baseline psychological state. Additionally, the Multidimensional Assessment of Interoceptive Awareness was excluded because it only strongly clustered with itself. Subsequently, baseline psychological state was represented using State-Trait Anxiety Inventory – Trait only, Anxiety Sensitivity Index, Mobility Inventory for Agoraphobia, UK Biobank Neuroticism Scale, and Epworth Sleepiness Scale. These questionnaires were all completed during the baseline test at sea level in Birmingham. Before clustering all variables were adjusted so that larger numbers represented a “worse” result and normalized via a z-transformation. The ability of the above variables to predict Lake Louise AMS score was further investigated using linear regression modelling.

## Results

### The Effect of Altitude on Metacognition

There was no significant difference in Mratio between visit 1 (140m) and visit 2 (2624m), HDI-0.2095, 0.6285. Additionally, there was no significant interaction effect (the effect of altitude) for the difference between visit 1 and 2 in the altitude group using control group as a comparison dataset (HDI, 0.5525, −0.5699). Neither was there a significant difference in the additional filter detection task variables between the two visits: perceptual sensitivity (RANOVA, F=0.5435, p=0.4660), perceptual bias (RANOVA, F=1.1436, p=0.2924), and average confidence (RANOVA, F=0.0001, p=0.9918).

### The Effect of a Daily Trek on Metacognition

There was no significant difference in Mratio between visit 2 (2624m) and visit 3 (2728m, after an 11-day trek), HDI −0.5203, 0.4295. Additionally, there was no significant interaction effect (the effect of daily exercise) between visit 2 and 3 in the altitude group using the control group as a comparison dataset (HDI, 1.6060, −0.1156). Neither was there a significant difference in the additional filter detection task variables between the two visits: perceptual sensitivity (RANOVA, F=0.8250, p=0.3701), perceptual bias (RANOVA, F=0.3104, p=0.5811), and average confidence (RANOVA, F=1.9743, p= 0.1691).

### Metacognitive Ability and AMS

The first regression model fitting Mratio, Lake Louise AMS score, and oxygen saturations from visit 2 (2624m) in the altitude group showed no significant association between metacognitive performance at that particular time point and corresponding Lake Louise AMS score (HDI, - 6.293, 0.2354) or oxygen saturations (HDI, −0.4579, 0.3530). The second regression model fitting Mratio from visit 1 in the altitude group and average Lake Louise AMS score, oxygen saturations, and heart rate from the highest camp on the expedition (4800m) found no association between metacognitive performance at baseline and measures of symptom burden at the highest camp on the expedition: Lake Louise AMS score (HDI, −0.1647, 0.5173), oxygen saturations (HDI, −0.2021, 0.4682), and heart rate (HDI, −0.3988, 0.1996).

### Questionnaires

The results of the self-report questionnaires from the baseline test of the control and altitude groups are shown in Table 1. There was a significant difference between two of the components of the Multidimensional Assessment of Interoceptive Awareness questionnaire: noticing and not worrying.

**Table 1.** Self-report questionnaire measures from baseline testing (in Birmingham, 140m) for the control and altitude group. An Anderson-Darling test was used to check whether data for each questionnaire was from a normal distribution reported as “True” for parametric data and “False” for non-parametric data; the two-groups were then compared using a two-tailed t-test or Wilcoxon rank sum test respectively. Uncorrected p-values are presented. A Bonferonni correction for multiple comparisons was used to adjust the level of significance to 0.0025; significant values are marked with an asterix (*). † MAIA – Multidimensional Assessment of Interoceptive Awareness.

| Questionnaire | Control Group<br>(median ± interquartile range) | Altitude Group<br>(median ± interquartile range) | Normally Distributed | p-value |
| --- | --- | --- | --- | --- |
| MAIA - Noticing | 3.6 ± 1.3 | 2.9 ± 1.3 | False | 0.1824 |
| MAIA - Not Distracting | 3.0 ± 1.3 | 1.8 ± 1.3 | False | < 0.001 * |
| MAIA - Not Worrying | 2.7 ± 1.0 | 3.8 ± 0.7 | False | < 0.001 * |
| MAIA - Attention Regulation | 3.0 ± 0.7 | 2.7 ± 1.6 | False | 0.4955 |
| MAIA - Emotional Awareness | 3.4 ± 1.4 | 3.0 ± 1.8 | False | 0.1196 |
| MAIA - Self-Regulation | 3.0 ± 1.8 | 2.4 ± 2.0 | False | 0.3919 |
| MAIA - Body Listening | 2.3 ± 1.3 | 2.2 ± 1.7 | False | 0.2020 |
| MAIA - Trusting | 4.0 ± 0.7 | 4.0 ± 1.0 | False | 0.5091 |
| Fatigue Severity Scale | 3.3 ± 1.2 | 2.9 ± 1.8 | False | 0.1208 |
| Epworth Sleepiness Scale | 7.5 ± 6.0 | 7.0 ± 3.0 | False | 0.6912 |
| State-Trait Anxiety Inventory - State | 33.5 ± 10.0 | 29.0 ± 14.0 | False | 0.2218 |
| State-Trait Anxiety Inventory - Trait | 38.0 ± 20.0 | 30.5 ± 8.0 | False | 0.0443 |
| Center for Epidemiological Studies Depression Scale | 11.5 ± 9.0 | 5.0 ± 6.0 | True | 0.0263 |
| Positive and Negative Affect Scale - Positive | 31.5 ± 10.0 | 39.0 ± 11.0 | False | 0.1130 |
| Positive and Negative Affect Scale - Negative | 14.5 ± 6.0 | 13.0 ± 6.0 | False | 0.4842 |
| Mobility Inventory for Agoraphobia - Alone | $1.4 \pm 0.4$ | $1.1 \pm 0.1$ | False | 0.0079 |
| Mobility Inventory for Agoraphobia - Accompanied | $1.2 \pm 0.3$ | $1.0 \pm 0.1$ | True | 0.0277 |
| Anxiety Sensitivity Index | $21.5 \pm 14.0$ | $10.5 \pm 5.0$ | False | 0.0028 |
| UK Biobank Neuroticism Scale | $2.5 \pm 6.0$ | $1.5 \pm 4$ | True | 0.2077 |
| Who Global Physical Activity Scale | $3900.0 \pm 4580.0$ | $3660.0 \pm 3040.0$ | False | 0.7879 |

### Baseline Psychology and AMS Symptoms

The results of the hierarchical cluster analysis are illustrated in Figure 2; the optimal number of clusters for this model was two. The first cluster was composed of mean oxygen saturation from the trek, Anxiety Sensitivity Index, and Epworth Sleepiness Scale. The second cluster included mean Lake Louise AMS score from the trek, Mobility Inventory for Agoraphobia – Alone and Accompanied, State-Trait Anxiety Inventory – Trait, and UK Biobank Neuroticism Scale.

**Figure 2.**
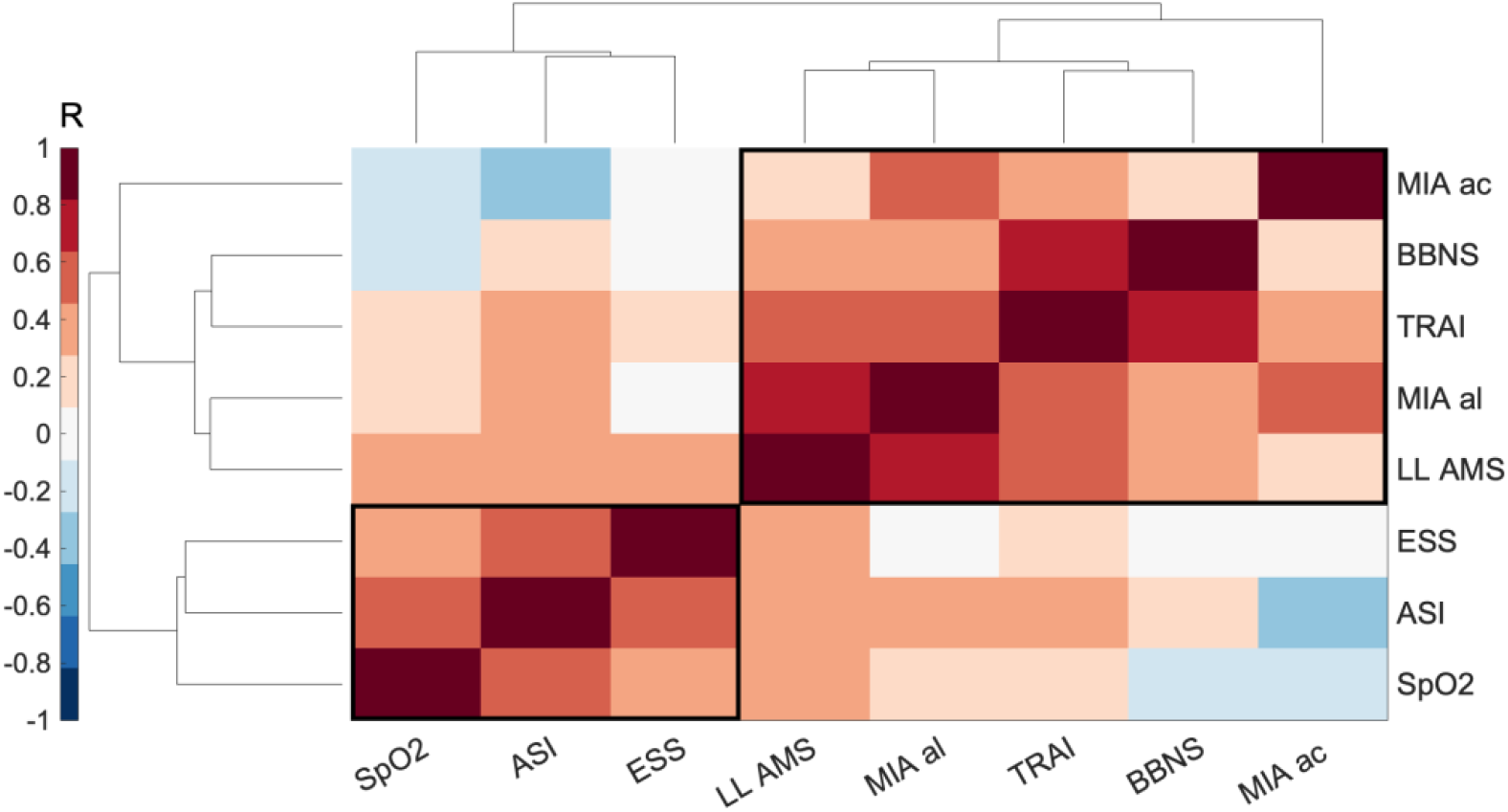
Clustergram: a correlation matrix of measures of Acute Mountain Sickness symptom burden over a high altitude trek and baseline psychological measures recorded at sea level prior to the expedition. Symptom burden was represented by mean Lake Louise Acute Mountain Sickness score (LL AMS) and mean oxygen saturations (SpO2) over the 11-day trek. Baseline psychology was represented by the following self-reported measures, State-Trait Anxiety Inventory – Trait only (TRAI), Anxiety Sensitivity Index (ASI), Mobility Inventory for Agoraphobia – alone (MIA al) and – accompanied (MIA ac), UK Biobank Neuroticism Scale (BBNS), and Epworth Sleepiness Scale (ESS). Strength of correlation is measured as a Pearson’s R-value displayed in the colour bar. The relationship between the groups of measures is demonstrated by the height of the dendrogram branches and the distance between neighbouring branches (in arbitrary units).

To explore whether any of these measures were predictive of the Lake Louise AMS score, they were fitted into a linear regression model: number of observations 16, degrees of freedom 8, root mean squared error 0.4546, adjusted r-squared 0.7926, F-statistic 9.1879, p=0.0028. The assumptions of linear regression, including normality of the residuals and homoscedasticity were met. In this model Epworth Sleepiness Scale (β_1_=0.5170, p=0.0314) and Mobility Inventory for Agoraphobia – Alone (β_1_=1.1574, p=0.0026) were predictive of Lake Louise AMS score. The other variables did not have a significant relationship with Lake Louise AMS score: mean oxygen saturation (β_1_=-0.0082, p=0.9750), State Trait Anxiety Inventory – Trait (β_1_=-0.0175, p=0.9432), Anxiety Sensitivity Index (β_1_=-0.3495, p=0.1750), Mobility Inventory for Agoraphobia – Accompanied (β_1_ =-0.6083, p= 0.0675), UK Biobank Neuroticism Scale (β_1_=0.0314, p=0.9064).

## Discussion

AMS is a poorly understood condition, in which symptoms are inadequately explained by measures of peripheral physiology. Interoceptive performance, as measured by the filter detection task, was not impaired on ascent to high altitude and did not demonstrate any relationship with AMS symptoms. However, average Lake Louise AMS score at high altitude clustered with self-reported measures of anxiety, agoraphobia, and neuroticism taken at sea level, but not average oxygen saturations at high altitude; this demonstrates the potential role individuals’ psychology has in explaining their AMS symptom burden.

There was no change in metacognitive performance between visit 1 at 140m and visit 2 at 2624m; therefore, ascent to altitude did not act as an “interoceptive challenge”. This may reflect the low incidence of AMS in our expedition group of predominantly experienced mountaineers at a moderate altitude of 2624m; the mean group Lake Louise AMS score at visit 2 was 0.83, with only one individual having a score of ≥3 classifying them as having mild AMS.

Neither was there a change in metacognitive performance between visit 2 and 3 after completion of an 11-day trek; therefore, daily exercise at altitude did not act as “interoceptive training”. Unfortunately, the trek was not as arduous as anticipated by the researchers. Additionally, the altitude group had a highly variable level of baseline activity level, with 74% of the group reporting >2000 minutes of metabolic equivalent time per week according to the Global Physical Activity Questionnaire (World Health Organization). Thus, it is possible that the trek did not present an adequate training stimulus to alter metacognition.

Metacognitive performance on arrival at altitude (visit 2) was not related to Lake Louise AMS score or oxygen saturations at that time point. Neither was baseline metacognitive performance (visit 1) predictive of AMS symptom burden (Lake Louise AMS score, oxygen saturations, or heart rate) at the highest point of the trek. It is possible that the filter detection task was not sufficiently sensitive to detect subtle changes in metacognition, particularly given the low prevalence of AMS on the expedition. Although, this same task has previously proven sensitive enough to detect interoceptive differences between asthma patient subgroups experiencing different levels of symptom severity (Harrison et al., 2020b).

Mood and physical fatigue appear to be important perceptual modulators impacting individuals’ symptom reporting (Harrison et al., 2020b; Finnegan et al., 2021), therefore we investigated the impact of these factors on AMS symptom burden in our cohort with hierarchical cluster analysis. Average Lake Louise AMS score over the trek clustered with self-reported measures of anxiety, agoraphobia, and neuroticism taken at sea level. Agoraphobia is a type of anxiety disorder where individuals fear being in circumstances where escape may be difficult (American Psychiatric Association, 2013). It has previously been linked to dysfunctional interoceptive processes, where individuals have increased perceptual sensitivity paired with a propensity to misconstrue bodily sensations as dangerous, leading to panic (Breuniger et al., 2017). Neuroticism has also been linked to interoceptive sensitivity (Pearson and Pfeifer, 2020) and certainly contributes to sensory perception in the Bayesian Brain, with neurotic individuals being predisposed to negative interpretations of sensations. Therefore, the clustering of Lake Louise AMS score with such psychological factors may represent a level of perceptual impairment in AMS prone individuals not detected by the filter detection task.

When fitted into a linear regression model, only two of the variables had a significant relationship with Lake Louise AMS score: Epworth Sleepiness Score and Mobility Inventory for Agoraphobia – Alone. Although a more specific symptom, sleepiness may be a manifestation of general fatigue, which is a theorised presentation of interoceptive dyshomeostasis (Stephan et al., 2016). However, this predictive relationship must be interpreted with caution given the sample size of this study and the number of variables included in the linear regression model. The authors suggest the limited inference that baseline psychological factors contributed to AMS symptom reporting in this study.

Notably, oxygen saturations did not cluster with Lake Louise AMS score and did not have a predictive relationship in the linear regression model. This suggests individuals’ psychological state may have contributed more to their experience of AMS symptoms on the trek than their oxygen saturations. Such a finding is unsurprising given that Lake Louise AMS is a general symptom score assessing headache, gastrointestinal symptoms, fatigue, dizziness, and ability to function with these symptoms. However, it is important to emphasise the impact of individuals’ psychology on Lake Louise AMS scores given its widespread use by researchers and commercial trekking agencies alike. We suggest that Lake Louise AMS scores should be interpreted with caution to prevent instances of serious pathology being missed.

### Limitations

The study had a small sample size, with only 18 participants in each group. A larger sample size would allow for exploratory factor analysis and stratification of participants into behavioural phenotypes.

We hypothesised that ascent to high altitude would impair interoceptive performance. However, the interoceptive tests took place at only moderate altitudes of 2624m and 2728m, inducing low levels of AMS in our study group. This was largely for pragmatic reasons, to allow testing to take place in an indoor environment, as the trekking group camped at higher altitudes.

## Conclusions

AMS remains a poorly understood condition, in which symptom burden is inadequately explained by measures of peripheral physiology. Interoceptive performance, as measured by the filter detection task, was not impaired in this study on ascent to altitude, or improved by daily exercise, and did not demonstrate any relationship with AMS symptoms. However, AMS symptoms were more closely related to self-reported psychological measures than oxygen saturations, demonstrating the contribution of psychological factors to individuals’ experience of AMS symptoms. Therefore, we advise caution in the interpretation of Lake Louise AMS scores by researchers and commercial trekking companies alike to ensure serious cases of AMS are not missed.

## Acknowledgements

We would like to thank all the members of the Birmingham Medical Expeditionary Society for facilitating this study on their 2019 Expedition to Sikkim.

## Author Confirmation Statement

BJT was involved in data collection, data analysis, and manuscript drafting.

CC, SJL, CL, and LM were involved in data collection.

OKH was involved in study design, data analysis, and manuscript drafting.

SLF was involved in data analysis and manuscript drafting.

SJEL was involved in study design and overview.

KP was involved in study conception, study design, data collection, data analysis, and manuscript drafting.

All authors were involved in final revision and approval of the manuscript.

## Author Disclosure Statement

The authors declare that the research was conducted in the absence of any commercial or financial relationships that could be construed as a potential conflict of interest.

## Funding Statement

This study was funded by JABBS Foundation.

## Supplemental Information

### Respiratory Filter Detection Task

We developed the respiratory filter detection task used in this study (Harrison *et al.*, 2020) by adapting the inspiratory resistance task used by Garfinkel et al., (2016a), to measure respiratory interoceptive ability. To interpret the results of this interoceptive test, its psychometric properties must first be fully understood, including whether there are learning effects on repeated measurement. Additionally, the minimum number of trials required to accurately determine metacognitive ability needs to be assessed to minimize the duration of the test, making it more applicable to clinical practice. Therefore we tested the following two hypotheses in this study: (i) there would be no learning effect on repeated completion of the filter detection task, and (ii) forty trials of the filter detection task would be equivalent to sixty trials in its calculation of metacognitive ability.

#### Learning Effect

To investigate whether there was a learning effect on repeated measures of the filter detection task, we used data from the control group, who completed the test three times without any “interoceptive training” or “challenges” between tests. The HMeta-d model was fit for each pair of tests (visit 1 and 2, visit 2 and 3, and visit 1 and 3), whereby individual participant data from each test is drawn from a multinomial distribution and the dependence caused by repeated measures within subjects is accounted for. To determine the significance of group differences in Mratio estimates of each of the test pairs, the HDI was calculated across the distribution of sample differences from each of the time points, as previously described for the Hmeta-d model (Fleming, 2017). A two-tailed 95% HDI that does not span zero determines a significant difference between the datasets.

There was no significant difference in metacognitive performance (Mratio) between visit 1 and 2 (HDI, −0.2040, 1.0173), visit 2 and 3 (HDI, −0.8549, 0.7159), or visit 1 and 3 (HDI, −0.1718, 1.0186). Neither was there a significant difference in any of the additional filter detection task variables between the three visits: perceptual sensitivity (number of filters), RANOVA, F = 2.74, p = 0.303; perceptual bias (bias towards yes or no), RANOVA, F = 2.62, p = 0.314; and metacognitive bias (average confidence), RANOVA, F = 0.277, p = 0.962.

This is the first study to have carried out repeated measures of the filter detection task (Harrison *et al.*, 2020). There was no learning effect on repeated completion of the task, validating its use in longitudinal studies. The control group alone was used to study this hypothesis so as to avoid the potential confounders of ascent to high altitude and daily trekking that occurred in the altitude group.

#### Forty Versus Sixty Trials of Filter Detection Task

We hypothesized that 40 trials of the filter detection task would be equivalent to 60 trials in calculating an individual’s metacognitive sensitivity (Mratio). Again, this hypothesis was tested using the control group, as their results have fewer confounding factors, remaining independent from the effect of ascent to altitude and daily trekking. Similar to hypothesis 4, the HMeta-d model was fit separately to each timepoint, using two sets of data within a multinomial model: 1) all available trials, and 2) only the first 40 trials of each participant. Then a HDI was calculated across the distribution of sample differences from each of the data comparisons described in hypothesis 1.

There was no significant difference in metacognitive performance as estimated using 40 trials versus 60 trials at visit 1 (HDI −0.4580, 0.3612), visit 2 (HDI −0.7707, 0.6866), nor visit 3 (HDI-0.5045, 0.8718). Similar to 60 trials, there was no significant difference in the additional filter detection task variables between the three visits using 40 trials: perceptual sensitivity (RANOVA, F = 2.74, p = 0.303); perceptual bias (RANOVA, F = 0.976, p = 0.630); and average confidence (RANOVA, F = 0.318, p = 0.944).

The perceived value of using only 40 trials rather than 60 trials was to reduce testing time, and therefore make it more amenable for integration into clinical practice. The control group was used to compare trial numbers to avoid the confounding features of the expedition to high altitude. However, throughout the course of the study we found the time difference between 60 and 40 trials to be minimal, with explaining the task and establishing the threshold number of filters to be the time-consuming stages of each test. As the time-saving is minimal and we know from model simulations that increasing the number of trials increases our confidence in the HMeta-d model (Fleming, 2017; Harrison *et al.*, 2020), we recommend continuing to use 60 rather than 40 trials in the filter detection task.

### Study Participants

The inclusion and exclusion criteria for participant recruitment are listed below.

#### Inclusion criteria

- All participants were willing and able to provide informed consent.
- All participant were adults aged 18 – 80 years.
- Only non-smokers were recruited.

#### Exclusion criteria

The participant did not enter the study if ANY of the following apply:

- Significant cardiac disease (e.g. heart failure, or pacemaker).
- Significant neurological disease (e.g. stroke, or neurodegenerative disease).
- Significant psychiatric disease (active treatment under psychiatric care).
- Significant metabolic disease (e.g. insulin dependent diabetes).
- Other respiratory diseases (e.g. sleep apnoea, chronic obstructive pulmonary disease, bronchiectasis, other restrictive lung disease, or severe asthma).
- Inadequate understanding of verbal and written information in English.
- Smoking history >20 pack years.
- History of prescription and non-prescription drug dependency (including alcoholism).
- Smokers.
- Travel above 1500 metres in the previous month.
- Travel across two or more time zones in the previous month.
- Travel across one time zone in the previous week.
- People who have worked night shifts in the previous month.

## Notes

### Competing Interest Statement

The authors have declared no competing interest.

